# Phase-Dependent Stimulation of the Hippocampus: A Computational Modeling Approach

**DOI:** 10.1101/2024.09.14.613022

**Authors:** Hsin-Pei Lee, Toren Arginteanu, Pawel Kudela, William S. Anderson, Yousef Salimpour

## Abstract

Phase-amplitude coupling (PAC) between brain oscillations of different frequencies plays a fundamental role in neural processing, and phase-dependent neuromodulation has emerged as a promising strategy to modulate PAC. In the hippocampus, theta-gamma PAC is critically involved in memory-related functions and information propagation. Computational models provide a valuable platform for investigating the neurobiological mechanisms underlying phase-dependent effects, bypassing the limitations of in vivo and in vitro experiments. In this study, we extended a previously published computational model of the hippocampal CA3 region using the NEURON and Python environments. A closed-loop autoregressive (AR) forward prediction model was employed to sample the network’s local field potential (LFP) in real time, enabling the precise calculation of phase-locked stimulus time points. Our results demonstrated the successful delivery of phase-locked current injections to all neuronal populations at both the peak and trough of theta oscillations. Phase-specific alterations in the theta band were observed during stimulation, along with enhanced theta-gamma coupling induced by peak-phase stimulation. Single neuron activity analysis highlighted the critical role of oriens lacunosum-moleculare (OLM) cells in modulating phase-dependent network dynamics. These findings underscore the potential of closed-loop stimulation systems to modulate PAC, with significant implications for the treatment of neurological disorders characterized by abnormal oscillatory activity, such as Alzheimer’s disease and other memory-related disorders.

**Significance Statement:** By employing a computational model of the hippocampal CA3 region, we reveal the ability of the phase-dependent stimulation technique to modulate phase-amplitude coupling, a critical mechanism in memory and information processing. Our findings highlight the importance of precise phase-locked stimulation and the key role of specific interneurons in regulating network dynamics. These insights lay the groundwork for developing targeted neuromodulation therapies to restore normal oscillatory patterns in the brain, with promising implications for treating memory-related neurological disorders.

## 1 Introduction

Brain oscillations, manifesting as local field potentials (LFP) and evident in electrocorticographic (ECoG) signals, arise from synchronized membrane potential changes in neuronal populations. Cross-frequency coupling (CFC) between brain oscillations of different rhythms reflects local cortical information processing and signal propagation across cortical regions. Analysis of ECoG signals from Parkinson’s disease and epileptic patients suggests CFC associations with motor and memory function (Salimpour and Anderson 2019). A specific type of CFC is phase-amplitude coupling (PAC), where the phase of the lower frequency oscillation modulates the amplitude of the higher frequency oscillation (Hyafil et al. 2015; Canolty and Knight 2010). Altered PAC has been implicated in various neurological disorders, including epilepsy, schizophrenia, obsessive-compulsive disorder (OCD), Parkinson’s disease (PD), and Alzheimer’s disease (AD) (Salimpour and Anderson 2019; de Hemptinne et al. 2013; Kondylis et al. 2016; R. Zhang et al. 2017; Goodman et al. 2018; Fujita et al. 2022). Notably, the modulation of the theta (4-9 Hz) oscillation phase on gamma (50-200 Hz) oscillation amplitude within the hippocampus plays a crucial role in memory functions (Salimpour and Anderson 2019; Vivekananda et al. 2021; Tamura et al. 2017). In patients with mild cognitive impairment and AD, the lower theta-gamma PAC correlates with lower memory performance (Musaeus et al. 2020). These findings emphasize PAC as a possible therapeutic target for treating neurological disorders.

Deep brain stimulation (DBS) has been commonly used to treat various neurological disorders, but the mechanisms remain unclear. It has been shown to reduce the abnormally elevated PAC in Parkinson’s disease (Swann et al. 2015; de Hemptinne et al. 2015). The current common stimulation paradigm applies a continuous, high-frequency (100–250 Hz) pulse train. However, there exist scarce randomized controlled clinical trials to test different stimulation paradigms, and the optimal parameters, including for treating epilepsy, remain unknown (Zangiabadi et al. 2019). Additionally, whereas beta oscillations have been identified as a biomarker in movement disorders, no single frequency oscillation band has been isolated for tracking seizures (Yang et al. 2023). Targeting PAC instead of a single frequency oscillation band may improve the current paradigm.

Despite the demonstrated implication of PAC in memory and disease, a comprehensive understanding of the underlying neurobiological mechanisms of phase-specific network effects in humans remains elusive (Neumann, Horn, and Kühn 2023). The exact method to target abnormal PAC to treat neurological disorders requires more understanding. Recent work by Duchet et al. aims to elucidate the optimal stimulation strategy to enhance or decrease PAC (Duchet and Bogacz 2024). Applications of phase-dependent stimulation have demonstrated effective modulation of cortical network activity and function in a phase-specific manner (Canolty and Knight 2010; Salimpour et al. 2022; Momi et al. 2022). Notably, Holt et al. demonstrated that continuous stimulation at the patient-specific phase of the beta oscillation in Parkinson’s patients could suppress the pathophysiological subthalamic nucleus (STN) beta oscillations (Holt et al. 2019).

In the brain-computer interface (BCI) application, Zhang et al. enhanced motor imagery-based performance with phase-dependent closed-loop stimulation (W. Zhang et al. 2021). Studies have also investigated phase-specific modulation of network activity at the cellular level. Peles et al. indicated synchronized neuronal firing at the preferred falling phase surrounding 168° of the beta band oscillations in monkeys (Peles et al. 2020). There are also studies to treat pathological PAC in the hippocampus (Bazzigaluppi et al. 2018). A mouse study showed the enhancement of spatial memory during either encoding or retrieval depending on the targeted theta phase (Siegle and Wilson 2014). Altogether, these studies suggest the promise of phase-dependent stimulation in neuromodulation and enhancing behavior.

Studying the network activity alterations caused by phase-dependent stimulation in humans poses several challenges. As such, computational modeling serves as a convenient surrogate. The NEURON simulation environment is a popular tool to model single neurons and networks of neurons (Carnevale and Hines 2006; Hines 2009). Existing hippocampal models built using NEURON have been validated to replicate experimental data and generate intrinsic theta-gamma modulation (Neymotin et al. 2013; 2011).

To investigate the underlying mechanisms of phase-dependent stimulation at the single-cell level, we extended the CA3 model by Neymotin et al. (Neymotin et al. 2013) incorporating phase-specific stimulation utilizing an autoregressive (AR) spectral estimation and time-series forward prediction method. The AR modeling has been presented to achieve phase-locking on theta oscillations (Chen et al. 2013). We hypothesized that stimulating different phases of the theta oscillation would result in different network effects. As such, we employed the algorithm to predict real-time phases of maximum separation: peak (0°) and trough (180°). The stimulation technique modeled direct stimulation of the neurons using depth electrodes. Our results demonstrated phase-specific effects on the network activity and highlighted the use of computational models to study the underlying neurobiological mechanisms of PAC.

## 2 Materials and Methods

### Stimulation Application

To observe phase-dependent effects *in silico*, we extended a published model of hippocampal CA3 (Neymotin et al. 2013). Simulations were run in the NEURON environment with a Python interpreter (Carnevale and Hines 2006; Hines 2009). The simulation environment allowed for randomization of synaptic inputs and network connectivity (Neymotin et al. 2013; 2011). Similarly, stimulation patterns can be customized and applied throughout the simulation. Simulation data was exported from Python and analyzed in Matlab (Mathworks, Inc.).

### Network Architecture

The cell types and connectivity were kept consistent with the original model (Neymotin et al. 2013). In total, the network consisted of 800 five-compartment pyramidal (PYR) cells, 200 one-compartment basket (BAS) interneurons, and 200 one-compartment oriens lacunosum-moleculare (OLM) interneurons (Figure 1A). The network parameters were tuned to generate theta (4-9 Hz), gamma (>30 Hz), and theta-modulated gamma oscillations (Canolty and Knight 2010). Additionally, current injections of 50 pA to pyramidal cells and -25 pA to OLM cells were delivered to provide a baseline activity as the isolated model did not allow for self- activation.

**Figure 1.**
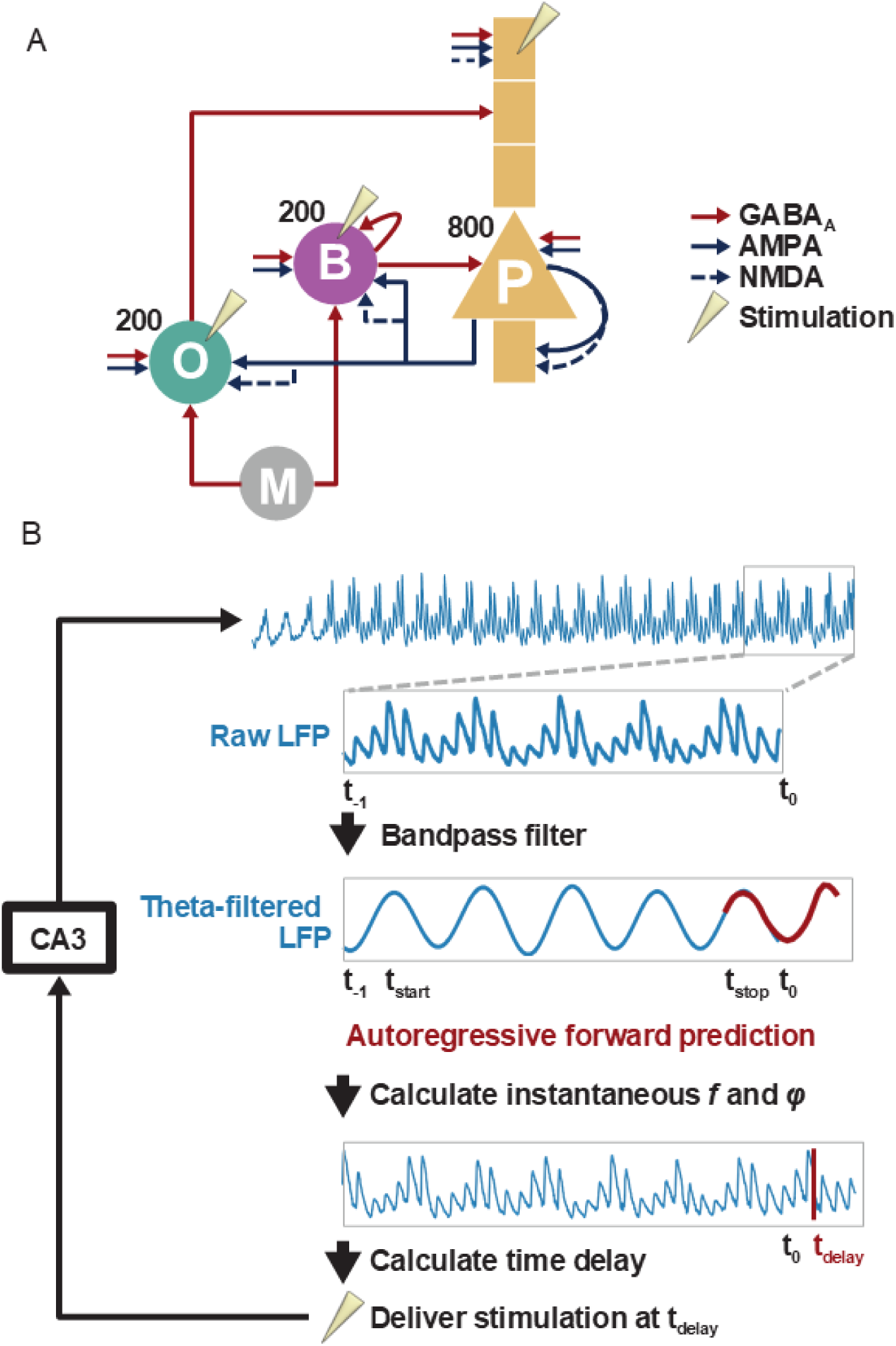
CA3 network architecture and phase-dependent stimulation: **A)** Schematic representation of the CA3 network architecture. P: 800 pyramidal cells; B: 200 basket cells; O: 200 OLM cells. M: external medial septal input to OLM and basket cells at theta frequency acts as a pacemaker. Arrows represent synaptic connections: inhibitory GABA_A_ inputs (red), excitatory AMPA inputs (solid blue), and excitatory NMDA inputs (dash blue). Cutoff lines represent external inputs. The needles represent the location of current injections to each cell population. **B)** Schematic representation of closed-loop phase-locked stimulation applied to the CA3 network. The raw LFP derived from the CA3 network is bandpass filtered within the theta band. The autoregressive forward prediction model then samples from the theta band oscillation to predict a segment of the theta band from t_stop_. Finally, the augmented theta band is used to calculate the time of the next stimulation pulse at the selected phase.

The voltage variables and current types of each cell were based on previously published models (Neymotin et al. 2013). All cells contained a leakage current, a transient sodium current, and a delayed rectifier current for action potential generation. Pyramidal cells contained a potassium type A current in all compartments for rapid inactivation. OLM cells contained a simple calcium-activated potassium current for long-lasting inactivation after bursting. In addition, all cells had a distributed hyperpolarization-activated current *I_h_* based on the HCN2 isoform parameterization. The *I_h_* currents modulated cell excitability and tuned the network’s theta frequency (Neymotin et al. 2013). To prevent compounding network effects with phase-lock stimulation, the *I_h_* currents were kept constant at the 1.0 scale setting for all cell types.

A total of 512,000 synapses were included in the model as described in the original publication (Neymotin et al. 2013). Pyramidal cells projected a mix of alpha-amino-3-hydroxy-5- methyl-4-isoxa-zolepropionic acid (AMPA) and N-methyl-D-aspartic acid (NMDA) synapses on basket interneurons and OLM interneurons. Basket interneurons projected gamma-aminobutyric acid A (GABA_A_) synapses on the soma compartments of pyramidal cells and OLM interneurons. OLM interneurons formed synapses on the distal dendrites of pyramidal cells via GABA_A_ receptors. AMPA and NMDA receptors had reversal potentials of 0 mV, and GABA_A_ receptors had reversal potentials of 280 mV. The cell connections were fixed throughout the simulations.

As in the original model, background activity simulated by excitatory and inhibitory synaptic inputs, following a Poisson process, was delivered to each cell to represent activities not explicitly modeled. Fast and slow background activities were generated by the activation of AMPA and GABAergic receptors at 1000 Hz and NMDA activations at 10 Hz, respectively. The model allowed for periodic inhibitory medial septal inputs to the interneurons at 10 Hz as a pacemaker to enhance the theta band of the network activity. We removed this input to focus on the intrinsic theta-band generation in the hippocampus. Inputs from the entorhinal cortex were modeled by slow excitatory activity at the last distal apical compartment of pyramidal cells to drive the sparse firing of pyramidal cells (Neymotin et al. 2013).

### Local field potential

According to the original Neymotin et al. model, the local field potential (LFP) was simulated by summing up the membrane potential of the most basal dendritic compartment subtracted by the membrane potential of the most apical dendritic compartment of all pyramidal cells (Neymotin et al. 2013; 2011). Since the pyramidal cells were arranged in a slice with the same orientation, the LFP calculation captured the network activity of the cells. The first and last 200 ms of the simulated LFP were removed before further signal analysis to avoid endpoint artifacts.

### Phase-lock Closed-loop Stimulation

Investigation of the phase-dependent-stimulation-induced effects required a consistent delivery of stimulation strictly locked to a phase of the theta band. We employed an autoregressive (AR) time-series prediction algorithm as described in Chen et al. (Chen et al. 2013). The AR prediction algorithm allowed a consistent prediction of when the next occurrence of a target phase angle would occur given an oscillation. First, the network LFP was calculated at a particular time point, *t_0_* (Figure 1B). Since we were interested in the modulation power of the theta (4-9 Hz) band, the raw LFP was filtered into the theta band using a zero-phase bandpass filter filtfilt() function from SciPy, a Python package. Next, we used the linear_model yule_walker() maximum likelihood estimation (MLE) function from the statsmodels.regression package in Python to forward predict a 300 msec segment of the theta band LFP. The prediction segment began at t_stop_ = t_0_ - 30 msec to eliminate the edge effects of bandpass filtering. The AR model sampled from t_start_ to t_stop_ where t_start_ = t_stop_ - 1500 msec and predicted with an order of 19. Additionally, the sampling rate of the theta band was downsampled from 10,000 Hz to 50 Hz. After augmenting the predicted segment to the original theta oscillation, the augmented oscillation was converted to an analytical signal using the signal.hilbert() function from SciPy.

The instantaneous phase and frequency of the analytical signal were used to calculate the next occurrence, t_delay_, of the target phase angle (Chen et al. 2013). Finally, a stimulus at t_delay_ was appended to the stimulation train. Each phase-lock stimulus was directly delivered as a current injection to each cell (PYR: third apical dendrite compartment; BAS: soma; OLM: soma) in the model. Since PYR cells were multi-compartmental, we tested the current injections to each compartment of the PYR cells. Current injections to the third apical dendrite compartment for all PYR cells achieved the best phase-lock result. Several amplitudes of current injections were tested. The amplitudes were kept small to match the scale of the model. We found an optimal phase prediction consistency with the stimulation amplitude at 20 nA for trough and -20 nA for peak when targeting both the peak and trough theta phases. A single pulse consisted of a constant amplitude, symmetrical biphasic stimulation of 0.2 msec duration.

We conducted two identical simulation experiments targeting the peak (0°) theta phase and the trough (180°) theta phase. Each experiment ran for 11.2 sec including, in sequential order, 3.2- sec pre-stimulation, 5-sec peri-stimulation, and 4-sec post-stimulation segments. The flexibility of computational simulation allowed us to pause the simulation periodically, employ the described AR prediction algorithm, append the predicted stimulus, and resume the simulation. The prediction algorithm was called every 90 msec after the occurrence of a stimulation. If no t_delay_ was predicted, the simulation advanced for 90 msec. A baseline experiment ran for 11.2 sec with no additional stimulation as control. The peristimulation segment of the baseline experiment was defined as the matching peristimulation time segment in peak and trough experiments. The network LFP data, stimulation patterns, and single neuron firing data were exported from Python for further analysis and visualization in Matlab.

### Band Power

The theta band power was calculated for the theta band of the network LFP across 1-sec windows with 50 msec overlaps and converted to decibels (dB). The gamma band power was calculated in the same manner for the gamma band of the network LFP.

### Spike Densities

The average cell firing densities were calculated by dividing the total number of spikes of each cell type in the 1 msec bin size by the total number of cells of the type and the duration of the window.

### Modulation Index

Phase-amplitude coupling (PAC) is a neural phenomenon where the phase of a low-frequency oscillation modulates the amplitude of a higher-frequency oscillation, playing a crucial role in cognitive processes. A common method for quantifying PAC is the Modulation Index (MI), introduced by Tort et al. (2010), which measures the degree to which amplitude fluctuations are organized by the phase of slower rhythms. To compute MI, the phase of a low-frequency band ( theta band) is divided into bins, and the mean amplitude of the high-frequency signal (gamma band) within each bin is computed. If the amplitude distribution across these bins shows non-uniformity, it indicates coupling, and the MI quantifies this deviation from a uniform distribution. This method has been widely applied in neuroscience research to study interactions between different neural oscillatory components during tasks like memory and attention (Salimpour et al. 2019, 2022)

### Stimulation artifact removal

In order to have an accurate PAC estimation using MI technique, we need to remove the fast transient change in the membrane potential which acts as an after stimulus artifact after each stimulation. To remove the artifact, we chose a standard duration for the artifact and the optimized AR model was utilized for predicting the modeling LFP during stimulation events. The predicted LFP was substituted for the stimulus artifact during active stimulation. This removed the transient stimulation artifact and provided accurate phase detection and PAC estimation. Since the electrical stimulation pulses are relatively narrow, the duration of the artifact is a limited interval during which the output of the AR model has a low prediction error with no discontinuity in the modeled LFP (Salimpour et al. 2022).

## 3 RESULTS

The goal of this study was to simulate a phase-lock stimulation targeting a specific phase of the hippocampal CA3 network’s theta band and observe phase-dependent effects on the network. Phase-lock stimulation was augmented on a previously published CA3 network. Theta phases of maximal difference across 180-degree separation have been suggested to modulate two opposing functional connectivity states in the hippocampus (Lurie et al. 2022). Therefore, we targeted two phases of 180-degree phase separation: theta peak at 0° and theta trough at 180° (Figure 2A and 2B). Raw network LFP was exported after each 11.2-sec simulation run and filtered into theta band using a zero-phase band-pass filter.

**Figure 2.**
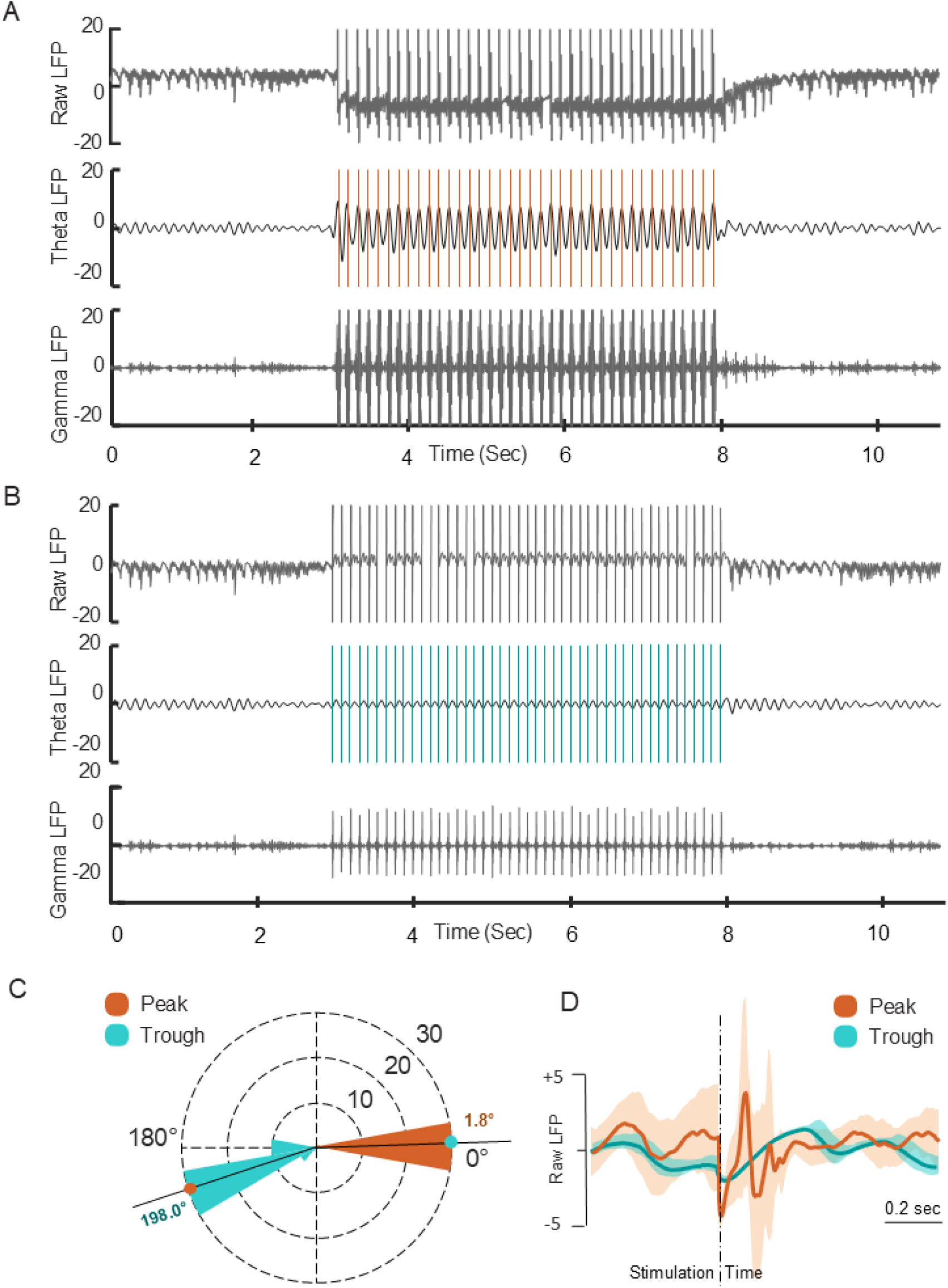
LFP model during stimulation: Network activity over time with peak (**A**) and trough stimulation phases. From top to bottom: raw (unfiltered) LFP, LFP theta band (4-9 Hz), LFP gamma band (50-200 Hz). The orange and blue vertical lines overlapping the theta band represent 20 nA current injections to each cell in the network in the peak and trough experiments, respectively. **C)** Phase plot of the corresponding phases of the stimulations in peak (*left*) and trough (*right*) experiments. A red dot indicates the median phase occurrence. **D)** LFP generated by the hippocampal model and isolated responses binned according to theta phase at stimulation delivery are shown. Phase-balanced grand average LFP in peak ( 0°) , through (180°), induced by current stimulation. Error bars indicate ±1 SD across LFP generation by hippocampal modeling.

The first step was to validate the robustness of the AR prediction algorithm. For each simulation run, the instantaneous phase was obtained by applying the Hilbert transform to the theta-filtered LFP. The phase of stimulation was the corresponding instantaneous phase at the time of stimulation. We observed highly consistent phases of stimulations in both peak (*N*=38,*M*=2.8°*,SD*=4.9°) and trough (*N*=45,*M*=198.8°*,SD*=10.6°) runs (Figure 2C). A total of 38 stimuli were applied in the peak experiment and 45 stimuli in the trough experiment.

### LFP amplitudes varied for theta peak versus trough stimulation

In a previous study on rodents, researchers found that the hippocampus is most receptive to input at specific theta-phase angles (Hasselmo et al., 2002). To see if this pattern holds true in a hippocampus model, we tested the responsiveness of the model to stimulation at different theta-phase angles. We estimated the theta phase at the start of each stimulation trial and grouped the trials into 45° intervals around the peak and trough phases. We used the stimulus artifact removal procedure (see the method session) to remove transient changes in membrane potential due to the current injection for stimulation. Then, we calculated the average peak and trough stimulation trials for the model’s generated LFP by taking the means within each interval (Figure 2D).

### Target Phase Altered Band Power

The theta band power was calculated for pre-, during-, and post-stimulation segments for each simulation run (Figure 3A). As expected from the constant setting of the model, the pre-stimulation band power was identical in all experiments.

**Figure 3.**
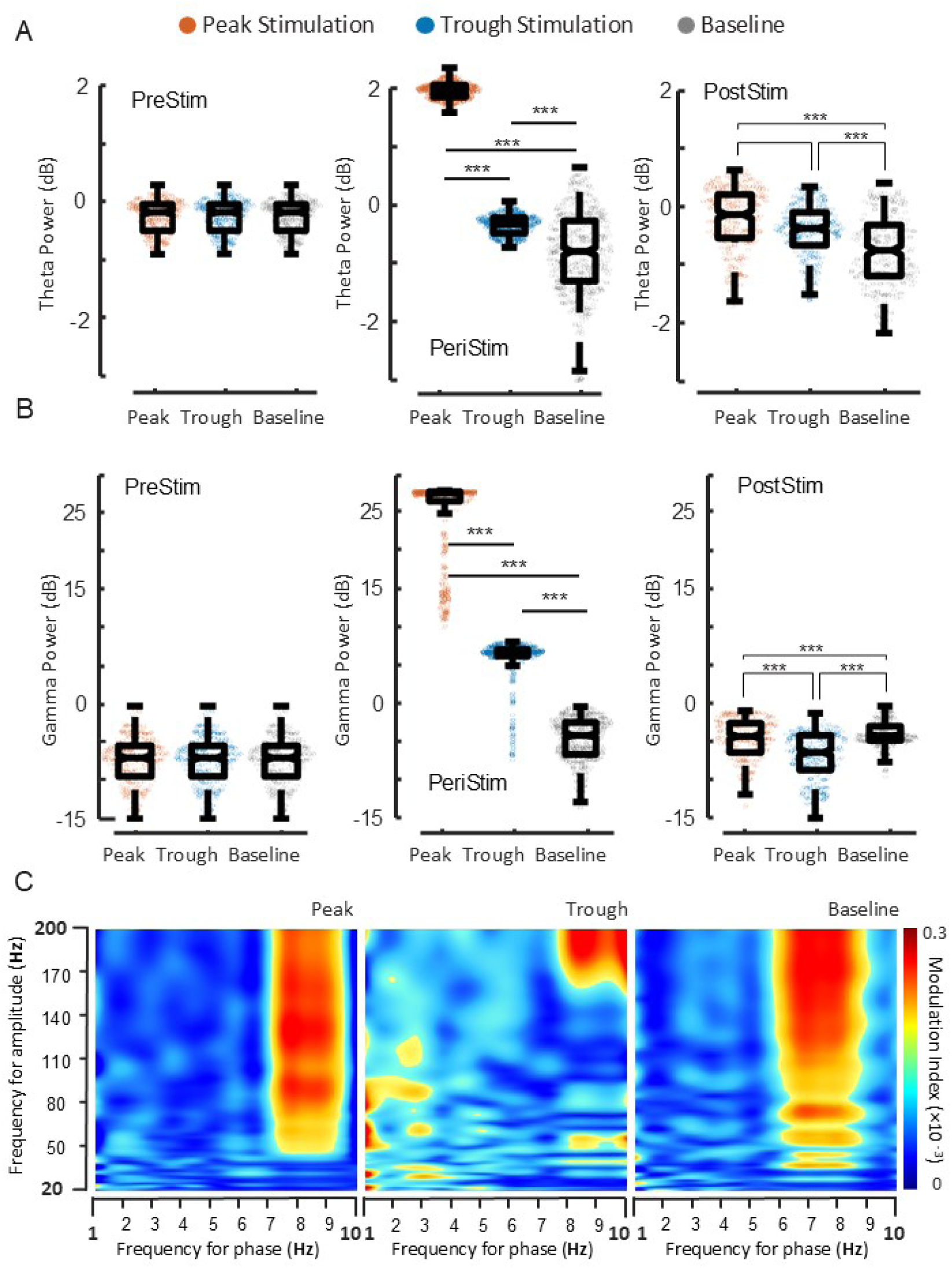
Theta, gamma, and theta-gamma coupling: **A)** Boxplot of theta power of the LFP oscillations over time calculated within the pre-stim, peristim, and post-stim segments. Plot shows comparison across peak (orange), trough (blue), and baseline (grey) experiments within each segment. **B)** Boxplot of gamma power across the same groups as in **A**; *** p<0.001. **C)** The level of coupling quantified by the modulation index between the theta frequency phase and gamma frequency amplitude for each experiment.

Peristimulation, the peak stimulations significantly elevated the theta band power compared to trough stimulations (p<0.001). Both peak and trough stimulations significantly elevated the theta band power from baseline (p<0.001). That said, the magnitude of change was greater for the peak simulation than the trough. The post-stimulation activity returned closer to the baseline level in both peak and trough simulations while maintaining trends observed peristimulation.

Changes in the gamma band power followed the same pattern as theta band power (Figure 3B). However, peristimulation, the magnitude of elevation in the gamma band powers was greater for both peak and trough stimulations than theta band power. Additionally, in the post-stimulation segment, the trough stimulations resulted in a significant decrease in gamma band power compared to baseline (p<0.001).

### Phase-dependent stimulations enhanced theta-gamma modulation

The theta-gamma modulation has been well established to be involved in spatial and long-term memory functions in humans (Tamura et al. 2017; Vivekananda et al. 2021). As such, the modulation index between the theta frequency phase and gamma frequency amplitude was calculated to investigate the impact of phase-lock stimulations on the level of coupling (Figure 3C). At baseline, the hippocampal CA3 network displayed theta band oscillations and theta-gamma modulation. The periodic MS input enhanced the intrinsic theta band power, which was associated with an increase in the gamma oscillation amplitude (Neymotin et al. 2013). With stimulations, both peak and trough stimulations shifted the theta frequency from 6-8 Hz to greater than 7 Hz. The coupling level increased with both stimulations. Nevertheless, the peak stimulations resulted in coupling at a wider range of frequencies for both gamma and theta oscillations. These results suggested a highly enhanced theta-gamma modulation associated with peak stimulations.

### Pyramidal activation leads increased theta band power while OLM activation leads reduced theta band power

To further understand the differential effects of peak and trough stimulations, we took a closer look at the single-cell level by taking advantage of the convenience of the NEURON environment to extract the membrane voltage and spiking activity of each cell (Figure 4A-D). We calculated the average firing rate of each cell population pre-, peri-, and post-stimulations. From the raster plots, the OLM population fired at wider intervals during peak stimulations compared to trough stimulations while the pyramidal and basket cells fired more frequently during peak stimulations. As shown in Figure 4E, the firing rates of pyramidal cells and basket interneurons elevated significantly during peak stimulations in comparison to trough stimulations (p<0.001). In contrast, the firing rate of the OLM population elevated slightly but not significantly during trough stimulations relative to peak stimulations. A slight elevation of OLM firing rate corresponded to increased pyramidal cell and basket cell firing rates. Our results agreed with the previous model that the OLM cells played a central role in modulating theta activity (Neymotin et al. 2013). Specifically, an increased OLM firing rate would dominate network inhibition and cause pyramidal and basket cells to decrease firing whereas a decreased OLM firing rate would result in higher firing rates of pyramidal cells that would feed forward to cause higher basket interneuron firings (Neymotin et al. 2013).

**Figure 4.**
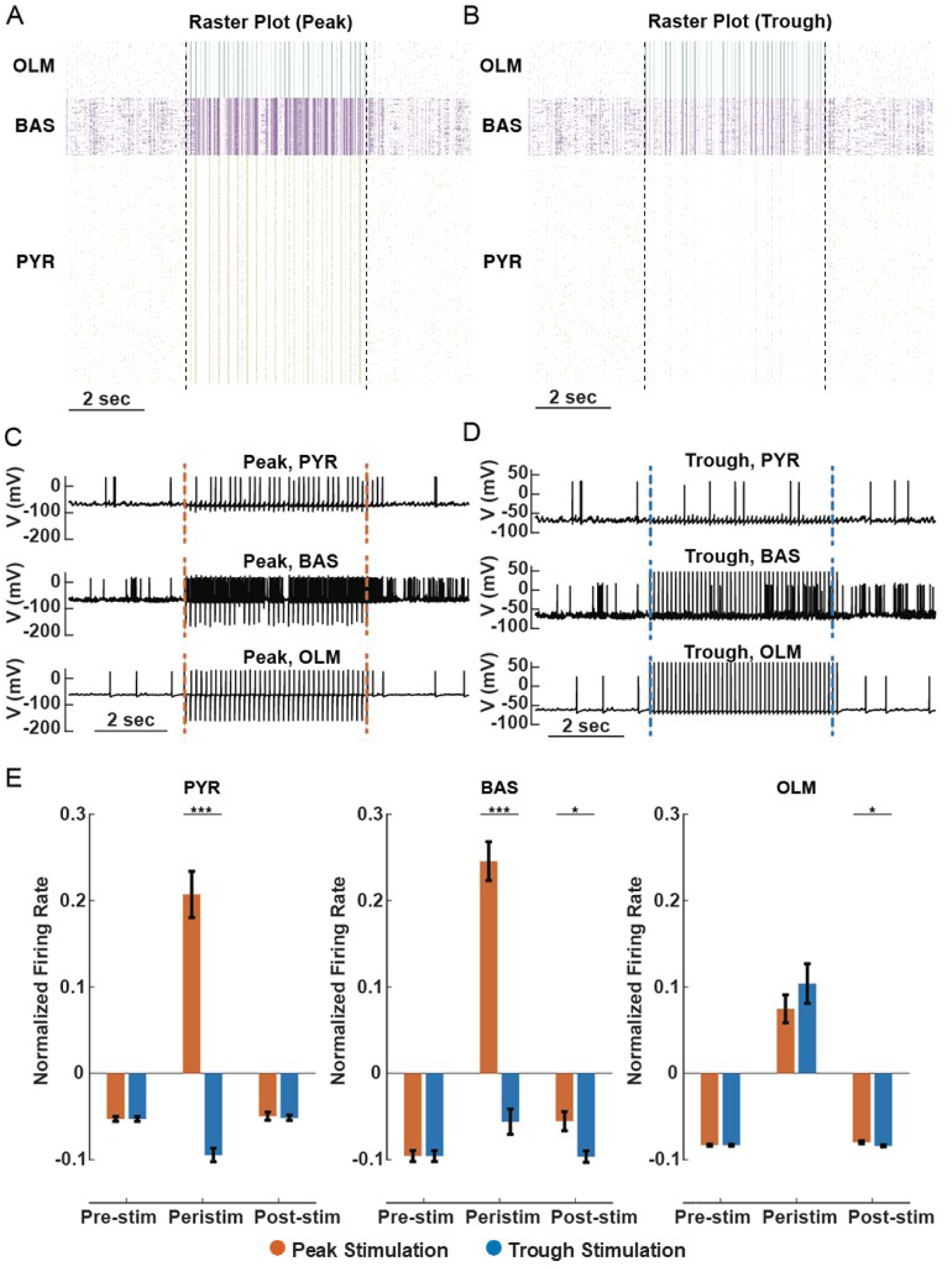
Single-cell response to the phase-dependent stimulation: A-B) Raster plot of single-cell spiking activity over time in peak (**A**) and trough (**B**) experiments. From top to bottom: OLM (green), BAS (purple), PYR (yellow). Dashed lines indicate the peristim segment. **C-D)** Membrane voltage over time of a representative cell in each cell population in peak (**C**) and trough (**D**) experiments. **E)** Bar plot comparing the normalized average firing rate of each cell population within the prestim, peristim, and post-stim segments across peak (orange) and trough (blue) experiments. ±SEM represented as error bars; * p<0.05, *** p<0.001.

### Comparison to clinical paradigm

We observed significantly higher firing rates of pyramidal and basket cells despite more stimuli in the trough experiment than in the peak experiment.

According to the concept of destructive interference (Kumari and Kouzani 2021), consecutive stimuli at opposing phases may cancel out phase-specific effects. As such, increasing the frequency of stimulation may be ineffective. Consistent phase-specific stimulations are potentially important to changing the network activity in a phase-specific manner. To demonstrate that phase-specific stimulations more effectively impact network activity than increased stimulation frequency, we applied an experiment mimicking the common clinical paradigm with continuous pulses at a high frequency of 130 Hz (Zangiabadi et al. 2019).

Analysis of the theta phase occurrence of the stimuli showed no phase-specificity across 650 stimuli (Figure 5A). The raster plot also indicated no observable pattern (Figure 5B). Comparing network activity, the theta and gamma band powers during the 130 Hz stimulation were more similar to baseline than peak and trough stimulations (Figure 5C and D). We then calculated the fold change of the firing rate of each cell population compared to baseline (Figure 5E). The fold changes of all three cell populations were nonsignificant in the 130 Hz experiment. Figure 5E shows that the firing rate of OLM increased significantly in both peak and trough experiments but not in 130 Hz experiment. OLM interneurons may play a crucial role in phase dependency. Overall, the 130 Hz experiment remained more similar to baseline activity at both network and cellular levels, suggesting the need for phase-specific stimulation.

**Figure 5.**
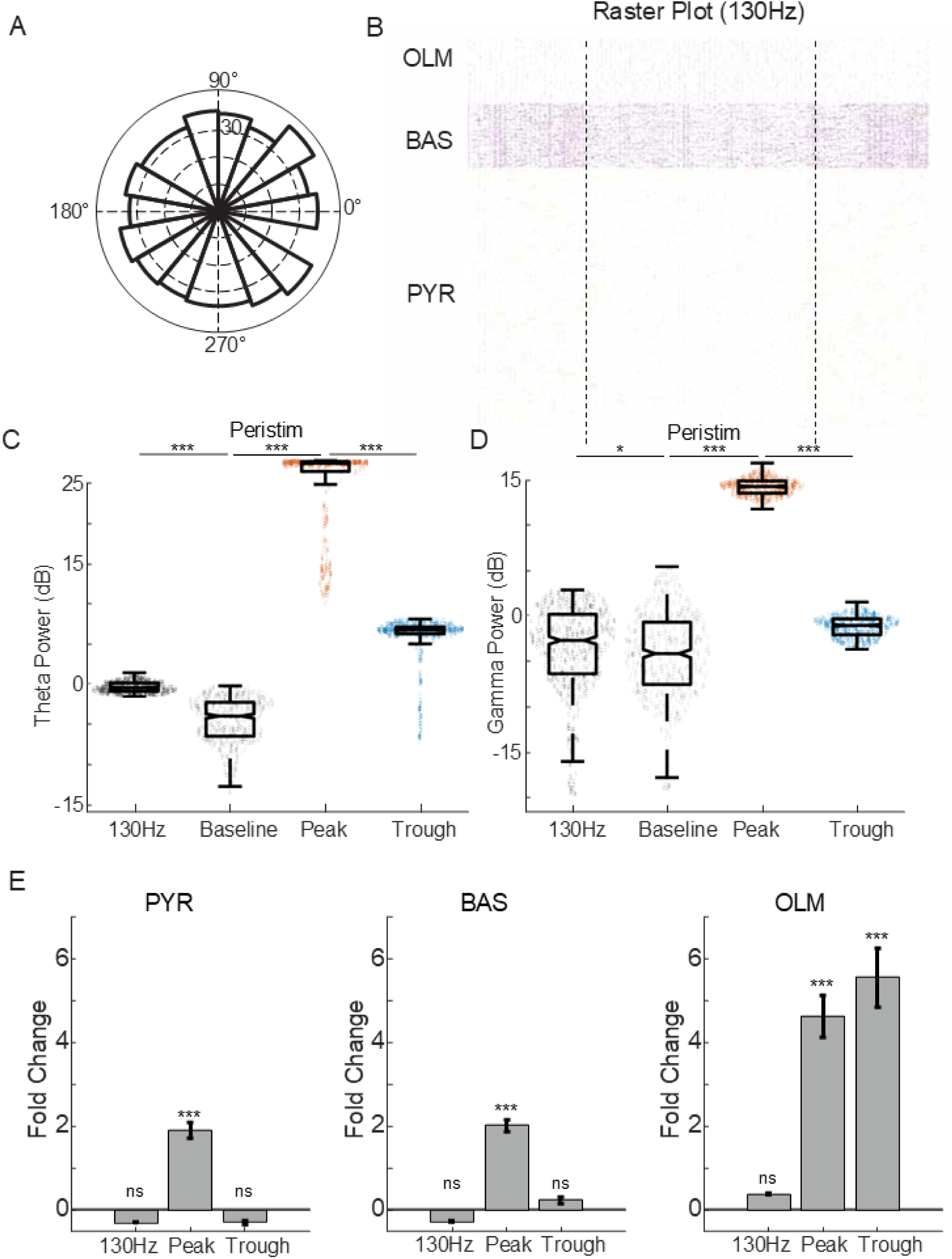
Single cell response to the open-loop stimulation: **A)** Phase plot of the corresponding phases of the stimulations in the 130 Hz experiment. **B)** Raster plot of single-cell spiking activity over time in 130 Hz experiment. From top to bottom: OLM (green), BAS (purple), PYR (yellow). Dashed lines indicate the peristim segment. **C-D)** Comparison of theta and gamma (**D**) band power across 130 Hz (dark grey), baseline (grey), peak (orange), and trough (blue) experiments during the peristim segment. **E)** Fold change of the mean firing rates of the three cell populations in 130 Hz, peak, and trough experiments compared to baseline. ±SEM represented as error bars; * p<0.05, *** p<0.001.

## 4 Discussion

We applied phase-lock stimulation targeting peak and trough theta phases to the CA3 hippocampal computational model using an AR prediction algorithm and observed phase-dependent network effects. A comparison of the theta band power during stimulations showed significant elevation with peak stimulations and significant depression with trough stimulations compared to pre-stimulation baseline activity. Both peak and trough stimulations increased the coupling level between the phase of theta oscillation and the amplitude of gamma oscillation. Additionally, peak stimulations resulted in a wider range of coupled frequencies in both theta and gamma rhythms. These results suggested the use of phase-dependent stimulation to modulate PAC.

The computational model allowed us to investigate the spiking activities of single cells. Firing rate analysis revealed a significant increase in pyramidal cells and basket cells only in the peak experiment. While insignificant, the OLM firing rates exhibited a decrease in firing rate in peak compared to trough experiments. Mirroring results of the original model, increased OLM firing rate could dominate network inhibition, feeding forward to decreased pyramidal and basket cell activities (Neymotin et al. 2013). Another study has also pointed out the importance of interneurons in the gamma synchronization (Tukker et al. 2007). These results underscored the important role of OLM in modulating network activity. The nonsignificant changes in firing rates across all cells in the 130 Hz simulation suggested that increasing the number of stimuli may not effectively change cellular activity.

Our results demonstrated the use of AR models to accurately predict the real-time theta oscillation phase and its validity as a technique for understanding differential network effects and modulating PAC. The model showed promise to mimic the *in vivo* environment, providing a platform for a more in-depth investigation into the neurobiological mechanisms underlying PAC at the single-cell level, not constrained to the hippocampal network. A recent study identified the PAC neuron population in regulating the hippocampal theta-gamma PAC associated with the cognitive control of working memory storage (Daume et al. 2024). Our computational model opens a convenient platform to more precisely investigate the role and properties of these neurons.

Although we demonstrated phase-dependent network and cellular activities, more work is required to understand the behavioral implications of phase-specific, closed loop stimulation. The theta-gamma PAC is associated with memory functions in the hippocampus (Axmacher et al. 2010; Lisman 2005; Madhavan et al. 2014; Mormann et al. 2005; Newman et al. 2013; Tort et al. 2009). A study comparing closed-loop and high frequency DBS on mice with parkinsonian symptoms shed light on the importance of tuning stimulation parameters based on the target structure (Cordon et al. 2018). In this study by Cordon et al., compared to closed-loop, the high frequency stimulation significantly reversed parkinsonian symptoms accompanied by reduction of beta oscillations. Such results highlight the importance of testing stimulation parameters with behavioral indications.

The model was limited in that it was small-scale, not human-based, and lacked intrinsic biophysical properties for long-term plasticity. As such, we were not able to observe a difference in the long-term post-stimulation effects. Several recent efforts have attempted to model PAC computationally (Ponzi, Dura-Bernal, and Migliore 2023; Duchet and Bogacz 2024; Vardalakis et al. 2024), suggesting a promise to extend our stimulation technique onto other models.

Additionally, phases other than peak and trough should be explored, including the rising and falling phases. Both the simplicity of the model architecture and the limited phase exploration may have contributed to our lack of observation in the reduction of theta and gamma band powers from baseline. Further study is required with more phase exploration and a more realistic, full-scale model that includes biophysical properties from human data and complex background inputs to the region of interest. Lastly, the results are limited *in silica* and have yet to be observed in human data.

To overcome clinical challenges and demonstrate phase-dependent effects, we implemented a phase-specific algorithm with an autoregressive forward-prediction algorithm. The model results revealed phase-specific alterations to the hippocampal network activity. These results suggested that phase-specific stimulation may be helpful in memory studies and in treating seizures in epileptic patients. Overall, this computational modeling serves as a pilot study and indicates a potential for clinical studies on phase-specific stimulation.

### Author contributions

**Hsin-Pei Lee:** Methodology, Software, Resources, Formal analysis, Validation, Writing original draft, Visualization, Data curation. **Toren Arginteanu**: Conceptualization, Methodology, Software, Review and editing. **Pawel Kudela:** Conceptualization, Investigation, Resources, Methodology. **William S. Anderson:** Project administration, Supervision, Conceptualization, Investigation, Resources, Methodology, Writing, Review and editing, Funding acquisition. **Yousef Salimpour:** Project administration, Supervision, Conceptualization, Investigation, Resources, Methodology, Writing, Review and editing, Funding acquisition.

### Disclosures

The authors have no actual or potential conflict of interest in relation to this study.

